# The viral protein NSP1 acts as a ribosome gatekeeper for shutting down host translation and fostering SARS-CoV-2 translation

**DOI:** 10.1101/2020.10.14.339515

**Authors:** Antonin Tidu, Aurélie Janvier, Laure Schaeffer, Piotr Sosnowski, Lauriane Kuhn, Philippe Hammann, Eric Westhof, Gilbert Eriani, Franck Martin

## Abstract

SARS-CoV-2 coronavirus is responsible for Covid-19 pandemic. In the early phase of infection, the single-strand positive RNA genome is translated into non-structural proteins (NSP). One of the first proteins produced during viral infection, NSP1, binds to the host ribosome and blocks the mRNA entry channel. This triggers translation inhibition of cellular translation. In spite of the presence of NSP1 on the ribosome, viral translation proceeds however. The molecular mechanism of the so-called viral evasion to NSP1 inhibition remains elusive. Here, we confirm that viral translation is maintained in the presence of NSP1. The evasion to NSP1-inhibition is mediated by the *cis*-acting RNA hairpin SL1 in the 5’UTR of SARS-CoV-2. NSP1-evasion can be transferred on a reporter transcript by SL1 transplantation. The apical part of SL1 is only required for viral translation. We show that NSP1 remains bound on the ribosome during viral translation. We suggest that the interaction between NSP1 and SL1 frees the mRNA accommodation channel while maintaining NSP1 bound to the ribosome. Thus, NSP1 acts as a ribosome gatekeeper, shutting down host translation or fostering SARS-CoV-2 translation depending on the presence of the SL1 5’UTR hairpin. SL1 is also present and necessary for translation of sub-genomic RNAs in the late phase of the infectious program. Consequently, therapeutic strategies targeting SL1 should affect viral translation at early and late stages of infection. Therefore, SL1 might be seen as a genuine ‘Achille heel’ of the virus.

## Introduction

The SARS-CoV-2, of the beta-coronavirus family, recently emerged as responsible for the Covid-19 world pandemic (Andersen et al., 2020; Zhou et al., 2020). Its genome is a positive single strand RNA molecule containing 29903 nucleotides entirely sequenced at the end of 2019 (Chan et al., 2020; Lu et al., 2020). The viral genomic RNA is capped at its 5’ end and polyadenylated at its 3’ end (Nakagawa et al., 2016). After entry in the infected cell, the viral genome hi-jacks the host translation machinery in order to produce the viral proteins required for the viral infectious program and the production of novel viral particles (Hartenian et al., 2020). Like many viruses, SARS-CoV-2 orchestrates viral translation concomitantly with the specific shut down of cellular mRNA translation. The goal of this silencing is dual. First, general cellular translation inhibition generates the large pool of ribosomes necessary to ensure efficient and massive synthesis of viral proteins. Interestingly, the cellular mRNAs coding for protein components of the translational machinery such as ribosomal proteins and translation factors are preserved from the overall translation inhibition presumably in order to maintain a functional translational machinery during viral translation (Rao et al., 2020). Secondly, the cellular translation silencing inhibits more specifically mRNA subsets that are involved in cellular immune responses to viral infection.

Viral translation begins with the expression from ORF1a that is translated into a polyprotein that is further processed by proteolytic cleavages to produce non-structural proteins (NSP) involved in the multiple steps of the general viral infectious program (Masters, 2006). The N-terminal proximal protein NSP1 is one of the first viral protein that is produced at the onset of the infectious program. NSP1 is required for efficient host cellular translation inhibition. First, NSP1 recruits a yet unidentified cellular endonuclease that promotes specific mRNA degradation on translated mRNAs (Kamitani et al., 2006, 2009). These cleavages occur on the ribosome during translation elongation. Importantly, viral mRNA transcripts are resistant to NSP1-mediated cleavages (Huang et al., 2011a). Secondly, NSP1 prevents translation initiation by interfering with the pre-initiation complex formation at multiple steps (Lokugamage et al., 2012a). NSP1 is thus directly responsible of general translation inhibition (Huang et al., 2011b; Lei et al., 2013; Narayanan et al., 2008; Tohya et al., 2009). However, its impact on translation is stronger on specific mRNAs. Among the targets of NSP1, mRNA subsets involved in specific cellular immune responses are primarily shut off. Thus, NSP1 suppresses type I interferon responses (Lei et al., 2020; Narayanan et al., 2008; Xia et al., 2020). The viral protein NSP1 binds to the host 40S ribosomal subunit with high affinity and a Kd in the nano-molar range (Lapointe et al., 2020). Its N-terminal domain was determined by NMR (Almeida et al., 2007), while the C-terminal domain of NSP1 contains an intrinsically disordered domain from residues 130 to 180 (Kumar et al., 2020). However, when bound to the ribosome, the NSP1 C-terminal domain is folded and binds tightly to the mRNA entry channel (Schubert et al., 2020; Thoms et al., 2020). NSP1 C-terminal residues from 148 to 180 interact with ribosomal proteins uS3 and uS5 and with helix h18 from the 18S rRNA (Schubert et al., 2020; Thoms et al., 2020). Interestingly, the NSP1 binding site overlaps with the binding sites of the initiation factors eIF1 and eIF3j (Lapointe et al., 2020) and, thus, the binding of NSP1 to the 40S prevents the formation of the 48S pre-initiation complex necessary for efficient translation (Brito Querido et al., 2020). NSP1 is also competing with the mRNA in the mRNA channel (Lapointe et al., 2020).

Recently, it has been shown that the N-terminal domain of NSP1 is required for viral translation by NSP1-bound ribosomes (Shi et al., 2020). The linker length between the N- and C-terminal domains is critical for viral translation (Shi et al., 2020). Beside the positive single strand genomic RNA, nine subgenomic RNAs (S, 3a, E, M, 6, 7a, 7b, 8, and N) are produced during the late phase of the infection by SARS-CoV-2 (Kim et al., 2020). All the viral transcripts are capped at their 5’ end with the first two nucleotides being ribose methylated and polyadenylated at their 3’ end (Lai and Stohlman, 1981; Yogo et al., 1977). The median length of the polyA tail on viral transcripts is 47 A residues (Kim et al., 2020). In coronaviruses, all viral transcripts contain the common so-called 5’ leader sequence (nucleotides 1 to 75) that forms the hairpins SL1, SL2, and SL3 (Kim et al., 2020; Miao et al., 2020; Sola et al., 2015).

Here we show that viral translation is evading NSP1-mediated inhibition because the viral transcripts, genomic and subgenomic RNAs, all contain a specific region of the leader sequence. More specifically, the sole hairpin SL1 is promoting NSP1 evasion by acting on NSP1 C-terminal domain in order to enable viral RNAs accommodation in the ribosome for their translation. The interaction between the SL1 RNA hairpin and the NSP1 C-terminal domain occurs while NSP1 remains bound on the ribosome. Therefore NSP1 acts as a ribosome gatekeeper to impair cellular translation and specifically promote viral translation.

## Results

### The evasion from NSP1 inhibition is due to *cis*-acting elements located in the SARS-CoV-2 5’UTR

To measure the impact of the SARS-CoV-2 5’UTR on viral translation, we inserted in a reporter construct the 5’UTR upstream of the *Renilla* luciferase coding sequence. As a control, a similar reporter containing the EMCV IRES was used. Using rabbit reticulocyte lysates (RRL), we measured translation efficiency of *Renilla* luciferase in absence and in presence of increasing concentrations of recombinant NSP1 (Figure 1). Translation efficiency of both constructs is reduced by NSP1. However, translation is significantly less affected with the SARS-CoV-2 5’UTR construct, indicating that the SARS-CoV2 5’UTR allows evasion from NSP1-mediated inhibition. This is in good agreement with previous studies that showed that NSP1 is indeed inhibiting EMCV-driven translation but not SARS-CoV-2 translation (Lokugamage et al., 2012b). To investigate further the evasion of SARS-CoV-2 viral translation from NSP1 inhibition, we analysed the formation of the ribosomal pre-initiation complex by fractionation on sucrose gradients. The RNA transcripts were radiolabelled at their 5’ ends and incubated in RRL. We used a minimal RNA containing the whole SARS-CoV-2 5’UTR and the first 12 codons of NSP1 coding sequence (nts 1-300) fused to a minimal portion of luciferase coding sequence (Figure 2). The formed pre-initiation complexes are then fractionated on sucrose gradient and collected in separate fractions. The presence of the radioactive RNAs in the collected fractions was then monitored by Cerenkov counting. With this experimental set-up, we detected the formation of 48S, 80S and disomes for EMCV and SARS-CoV-2 constructs (Figure 2A). In the presence of 0.8 μM NSP1, the formation of these complexes is drastically reduced with the EMCV transcript. In contrast, the SARS-CoV-2 5’UTR transcript allows for the formation of 80S complexes to the same extent and with only a slight reduction of disomes. This experiment confirmed that the evasion from NSP1 inhibition is due to *cis*-acting elements located in the SARS-CoV-2 5’UTR.

**Figure 1:**
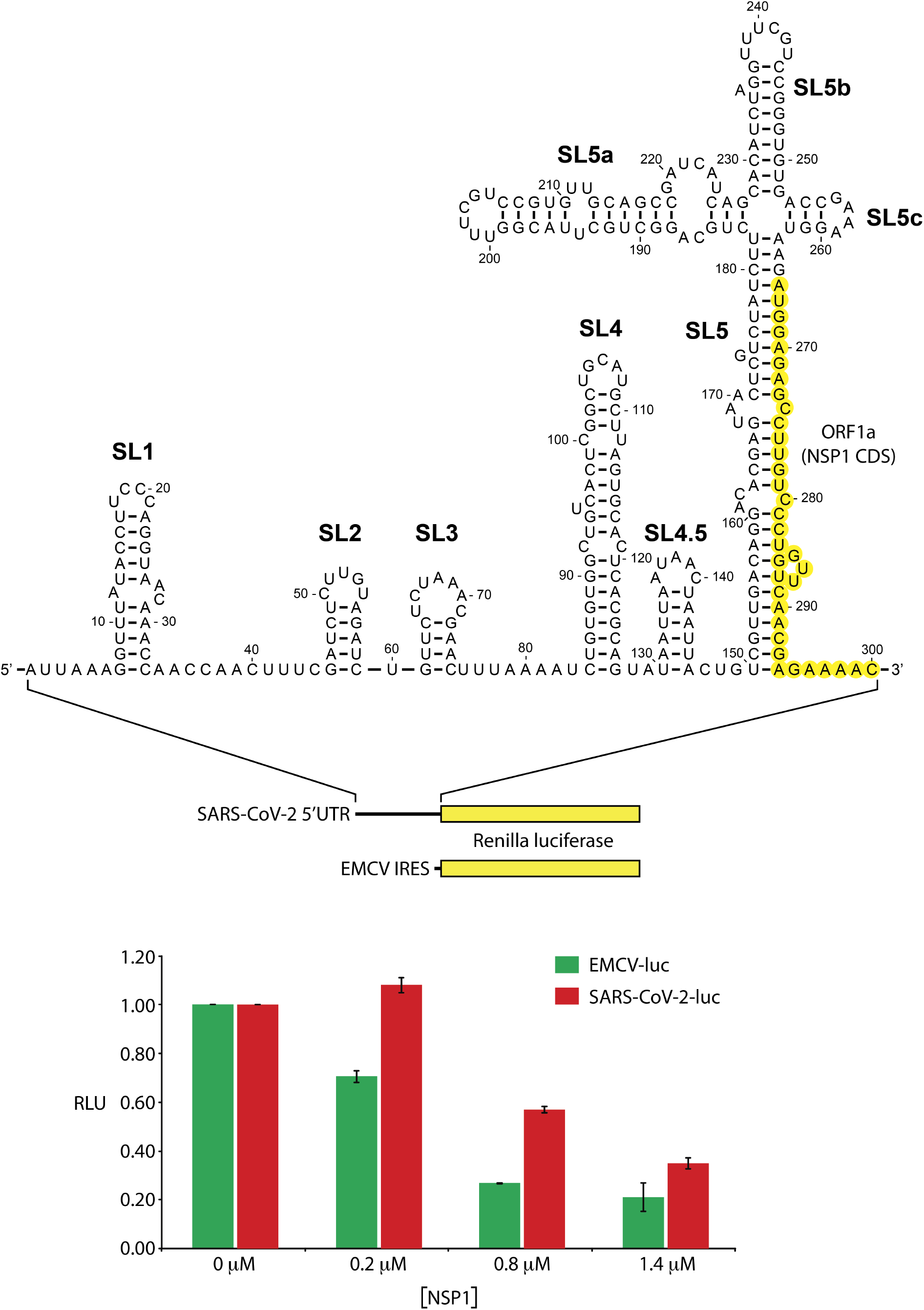
Translation inhibition by viral NSP1. The first reporter construct contains the SARS-CoV2 5’UTR plus the 12 N-t codons of NSP1 (nucleotides 1-300) fused to *Renilla* luciferase coding sequence. The second reporter contains the EMCV IRES upstream of the *Renilla* luciferase coding sequence. Translation efficiency was measured in absence or in presence of 0.2, 0.8 and 1.4 μM of recombinant NSP1. The average relative activities from three independent experiments are represented in the histogram.

**Figure 2:**
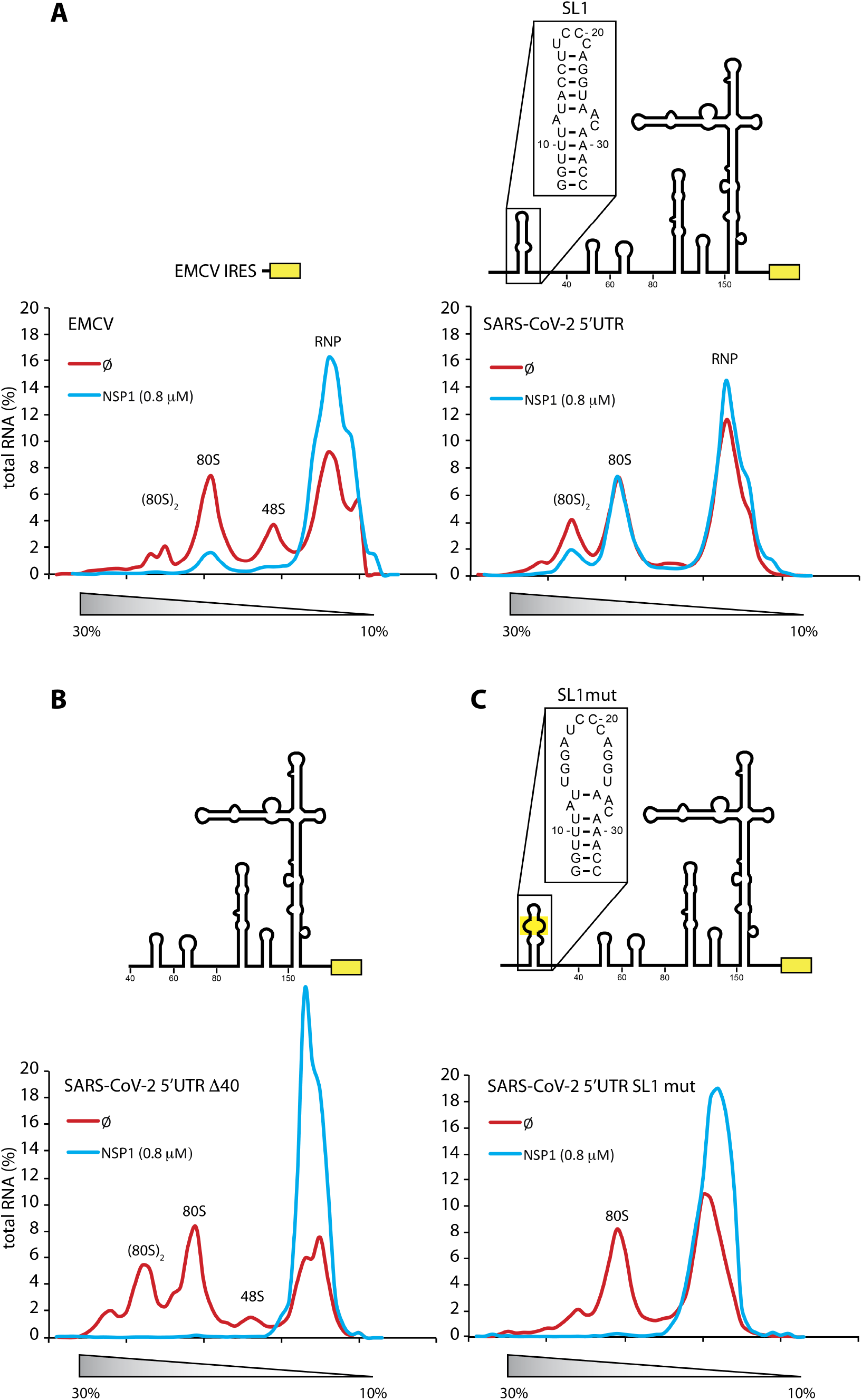
SL1 is required in the SARS-CoV-2 5’UTR for NSP1 evasion. Pre-initiation complex formation analysis on 10-30% sucrose gradients using radiolabelled RNA at their 5’ end. The SARS-CoV-2 RNA transcript was radiolabeled with a radioactive m7G cap at its 5’ end. The pre-initiation complexes were fractionated on 10-30 % sucrose gradients. The presence of radioactive RNA was monitor by Cerenkov counting of all the fractions. The plots represent the percentage of radioactive RNA that was used for complex formation. The pre-initiation complexes were formed in the absence (red line) or in presence of 0.8 μM of recombinant viral NSP1 (blue line). The positions of ribonucleoproteins (RNP), the 48S, 80S and disomes are indicated above the curves. (A) Translation initiation complexes analysis on sucrose gradients with EMCV reporter mRNA (left panel) and with SARS-CoV-2 reporter mRNA (right panel). Translation initiation complexes analysis on sucrose gradients with a truncated SARS-CoV-2 (B) or a SARS-CoV-2 5’UTR containing a mutation that disrupts SL1 (C).

### The apical part of SL1 is absolutely required for NSP1 evasion

In order to identify precisely the *cis*-acting elements, we repeated the experiments with truncated SARS-CoV-2 5’UTRs. With the 5’ proximal 40 nucleotides (Δ40) deleted, the protection towards NSP1-mediated inhibition is totally abrogated, which indicates that this part of the 5’UTR contains essential *cis*-acting elements (Figure 2B, Supplemental Figure 1). This region of the 5’UTR contains the predicted hairpin SL1 (Rangan et al., 2020) that was confirmed by probing experiments (Miao et al., 2020). Next, we introduced four mutations in the upper part of SL1. These mutations prevent the formation of the apical stem-loop of SL1 (called SL1 mut) (Figure 2C). We have verified that the introduced mutations do induce the opening of SL1 (Supplemental Figure 2). Again, the formation of pre-initiation complexes is totally prevented with SL1 mut, indicating that evasion to NSP1-mediated inhibition is abrogated. Thus, the apical part of SL1 is absolutely required for NSP1 evasion.

### A fully functional NSP1 is necessary for translation inhibition

Mutations of residues K164 and H165 of NSP1 into alanines totally abolish the binding to the 40S subunit (Kamitani et al., 2009). Likewise, the mutations R124A and K125A inhibit NSP1 ability to promote translation inhibition (Lokugamage et al., 2012a). We tested the effect of these mutations in NSP1 on the pre-initiation complex formation with the SARS-CoV-2 containing a mutated SL1. As expected, the wildtype NSP1 severely reduces the formation of the pre-initiation complexes when SL1 is mutated. In contrast, none of the two NSP1 mutants affect the translation indicating that NSP1 has to be bound to the ribosome to efficiently inhibit translation (Supplemental Figure 3). Altogether, these experiments demonstrate that the presence of the *cis*-acting element SL1 and a NSP1 protein able to bind ribosomes are necessary to promote a viral translation resistant to NSP1-mediated inhibition.

### The apical part of the SL1 RNA hairpin is solely responsible for NSP1 resistance

Next, we tested if SL1 is solely responsible for NSP1-resistance. For that purpose, we measured the impact of NSP1 on translation of another reporter construct containing the β-globin 5’UTR upstream of the renilla coding sequence. Translation is significantly inhibited in the presence of increasing concentrations of NSP1, confirming the general inhibitory effect of NSP1 bound on the ribosome (Figure 3A). We then transplanted the 5’ proximal 40 nucleotides of the SARS-CoV-2 5’UTR, which contain the SL1 hairpin, upstream of the β-globin 5’UTR upstream. The sole presence of SL1 allowed a significant protection against NSP1 inhibition. In contrast, when SL1 mut is transplanted, no protection is observed. Since the sub-genomic RNAs contain the so-called leader sequence that encompasses SL1, SL2 and SL2 in their 5’UTR, we checked whether SL2 and SL3 are also required for NSP1 evasion. Indeed, addition of SL2 and SL3 only slightly improves the translation efficiency in the presence of NSP1 (Figure 3B). This experiment indicates that sub-genomic RNAs can also evade to NSP1-mediated inhibition because they harbour SL1 in their 5’UTR. These data confirm that the apical part of SL1 is essential for NSP1 evasion. Altogether, these results indicate that the evasion of the SARS-CoV-2 RNAs to NSP1 inhibition is due solely to the presence of SL1 in the 5’UTR leader sequence.

**Figure 3:**
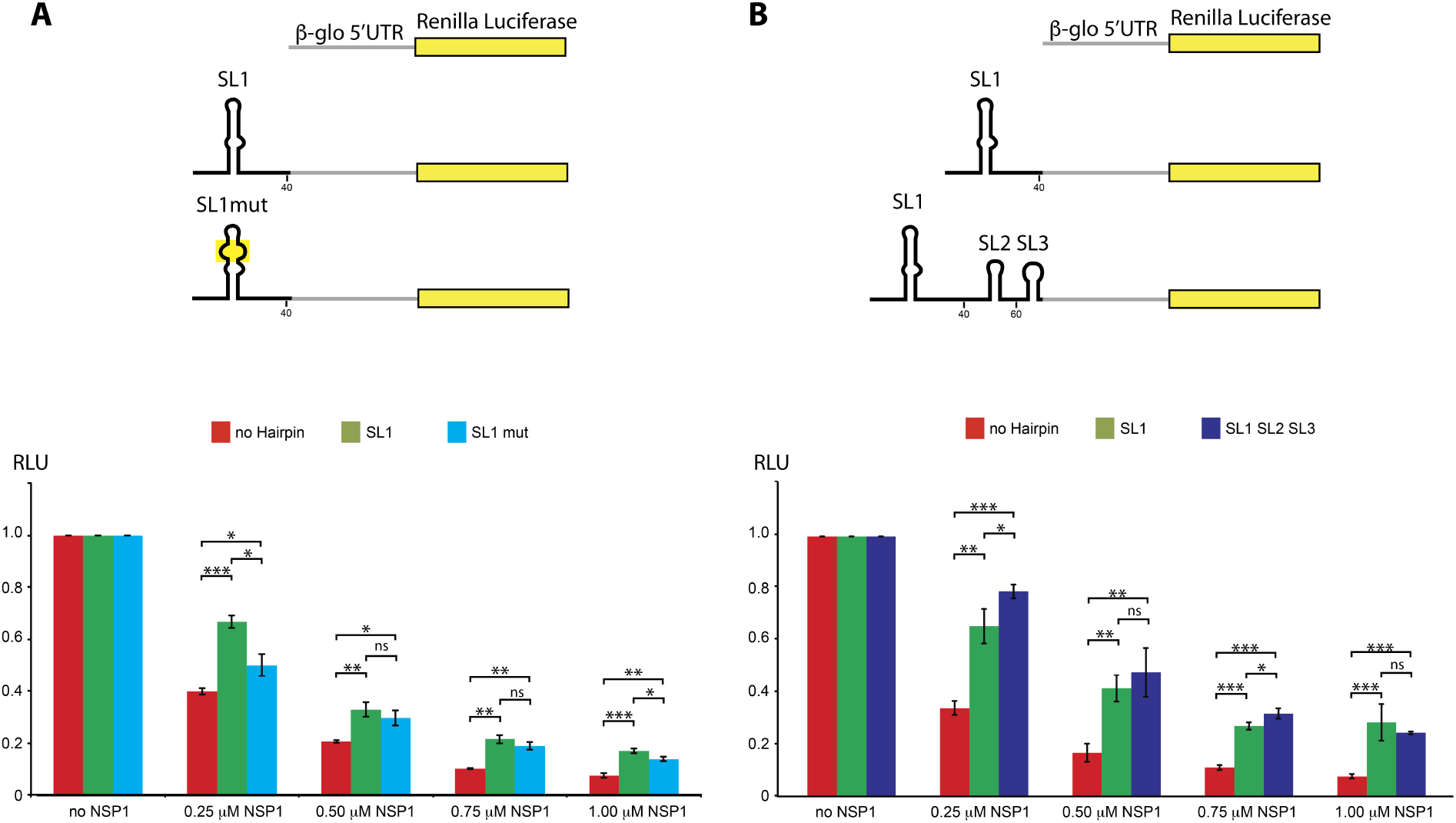
SL1 is sufficient to confer resistance to NSP1 inhibition. (A) Luciferase reporter mRNAs containing β-globin 5’UTR, SL1-β-globin 5’UTR and SL1mut-β-globin 5’UTR were used to measure their translation efficiency in RRL in the absence or in the presence of 0.25, 0.50, 0.75 and 1 μM of recombinant NSP1. The average relative activities from three independent experiments are represented in the histogram. The activity of the reporter β-globin 5’UTR luciferase in the absence of NSP1 is used as a control for normalization. (B) Luciferase reporters containing β-globin 5’UTR or with SL1, or with SL1-SL2-SL3 in their 5’UTR. Standard deviations or translational activity for each transcript are shown and calculated from three independent experiments. ns: non significant; *: 0,05 < p value < 0,01; **: 0,01 < p value < 0,001; ***: 0,001 < p value; based on Student’s t-test.

### Free NSP1 has no affinity for RNA

A putative mechanism would be that SL1 directly interacts with NSP1 and thereby removes NSP1 from the ribosome to allow access to the mRNA channel of the ribosome during the initiation process of mRNA accommodation. To evaluate such a mechanism, we tested the RNA binding ability of NSP1 (Figure 4A). We used radiolabelled RNAs from SARS-CoV-5’UTR with EMVC and HCV IRES as negative controls. None of the RNAs tested are bound by NSP1 even when 20 μM of NSP1 was used for Electrophoretic Mobility Shift Assay (EMSA). We concluded from these experiments that free NSP1 has no affinity for RNA even when it contains SL1 like the SARS-CoV-2 5’UTR.

**Figure 4:**
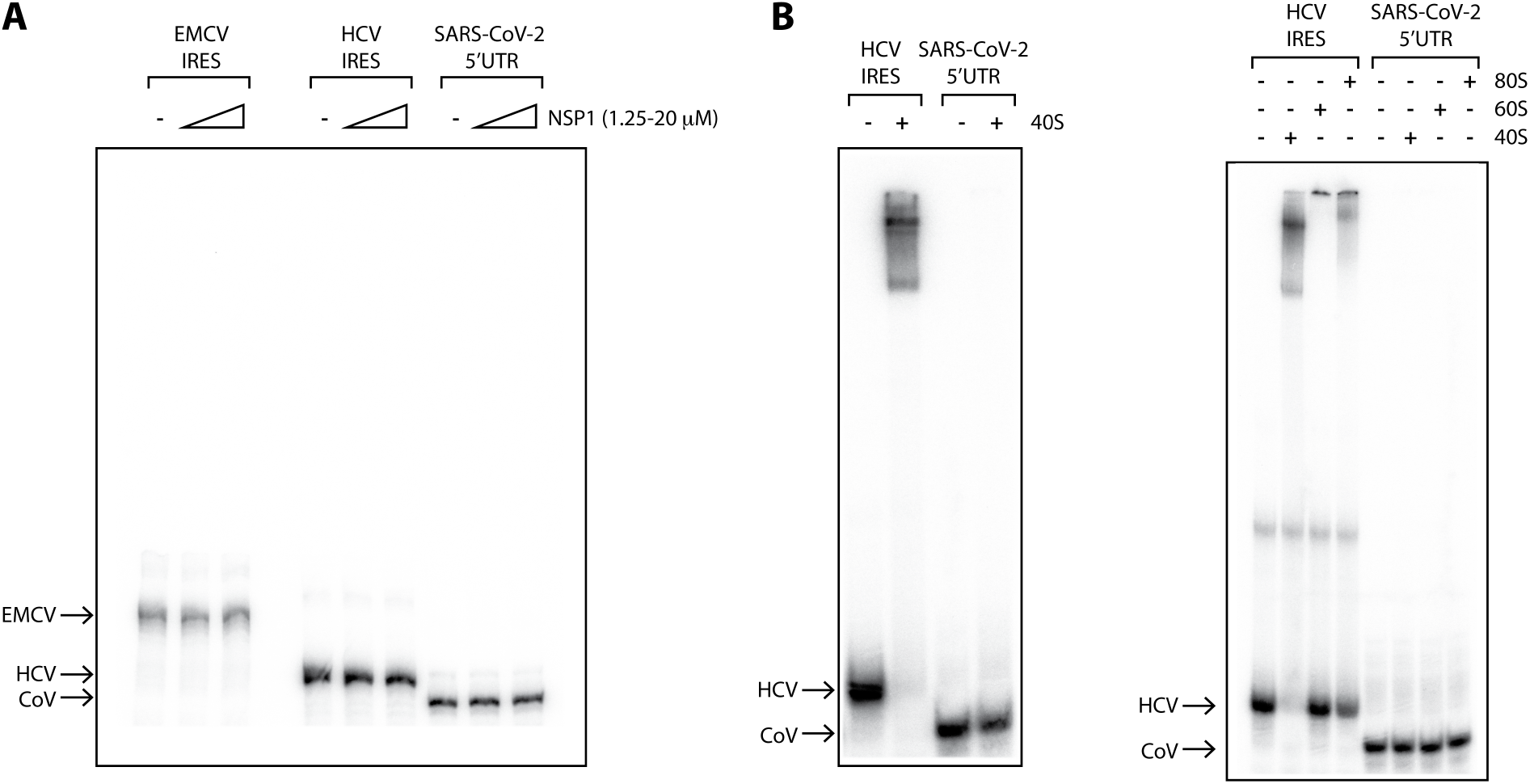
NSP1 has no RNA binding ability on its own. (A) Electro Mobility Shit Assay using radiolabelled RNA containing EMCV IRES, HCV IRES and the SARS-CoV-2 5’UTR. The RNAs were incubated in the presence of 1.25 to 20 μM recombinant NSP1 and loaded on native polyacrylamide gel. (B) The SARS-CoV-2 5’UTR was also used to test its binding to ribosomal 40S (left panel) and to 40S, 60S and 80S (right panel). The HCV IRES was used as a positive control. The positions of the free RNA are indicated by arrows.

### The SARS-CoV-2 5’UTR does not bind to any component of the ribosome on its own

However, NSP1 has a strong affinity for the 40S ribosomal subunit and its C-terminal domain binds into the mRNA channel with a Kd in the nanomolar range (Lapointe et al., 2020). We therefore tested whether a SARS-CoV-2 transcript is able to bind to pure ribosomal subunits by EMSA. To validate our assay, we used the HCV IRES as a positive control, since it was shown earlier that it interacts specifically with the 40S ribosomal subunit (Filbin et al., 2013; Fuchs et al., 2015; Quade et al., 2015; Yamamoto et al., 2015) and even with full 80S (Yokoyama et al., 2019). As expected, the HCV IRES interacts with purified human 40S ribosomal subunit and with the 40S subunit from the complete 80S ribosome. In contrast to the HCV IRES, the SARS-CoV-2 5’UTR is not able to bind the 40S, as well as the 60S or the 80S particles (Figure 4B). In summary, NSP1 has a strong affinity for ribosomal 40S subunit but the SARS-CoV-2 5’UTR cannot bind to any component of the ribosome on its own.

### The SARS-CoV-2 5’UTR promotes the assembly of pre-initiation complexes in presence of NSP1: the interaction between NSP1 and SL1 frees the mRNA accommodation channel while maintaining NSP1 bound to the ribosome

But our results also indicate that the SARS-CoV-2 5’UTR promotes the assembly of pre-initiation complexes in the presence of NSP1. In order to determine whether NSP1 is removed from the assembled pre-initiation complexes, we used a previously established protocol that yields purified pre-initiation complexes programmed with SARS-CoV-2 5’UTR (Chicher et al., 2015; Martin et al., 2016; Prongidi-Fix et al., 2013). The principle is to use a chimeric molecule composed on one hand by the RNA region encompassing the SARS-CoV-2 5’UTR followed by a small coding sequence and a DNA oligonucleotide coupled to Biotin at its 3’ end (Figure 5A). The hybrid molecules are then immobilised on magnetic streptavidin beads and incubated in RRL in the presence of cycloheximide to allow the formation of pre-initiation complexes. The complexes are then eluted by DNase digestion that removes the Biotin and the DNA linker. The composition of the eluted ribosomal complexes is determined by mass spectrometry analysis. We performed in parallel two experiments with the SARS-CoV-2 5’UTR in the presence or in the absence of NSP1. Each experiment was repeated three times (Figure 5B). In both experiments, pre-initiation complexes were efficiently purified as attested by the presence of 40S and 60S ribosomal proteins and eukaryotic initiation factors (eIF) (Figure 5C). We found the full set of ribosomal proteins from the 60S and the 40S ribosomal subunits (50 proteins and 36 proteins respectively) (supplementary table 2). Interestingly, we found that NSP1 is still present in the pre-initiation complexes formed in the presence of NSP1. This indicates that the purified pre-initiation complexes contain NSP1 still bound on the ribosome. Thus, these experiments imply that the mRNA channel is accessible to the mRNA and that NSP1 C-terminal domain must have been remodelled in order to allow mRNA accommodation. Since NSP1 is present in these pre-initiation complexes, this suggests that NSP1 remains attached to the ribosomal subunit with its N-terminal domain (according to data from (Shi et al., 2020)).

**Figure 5:**
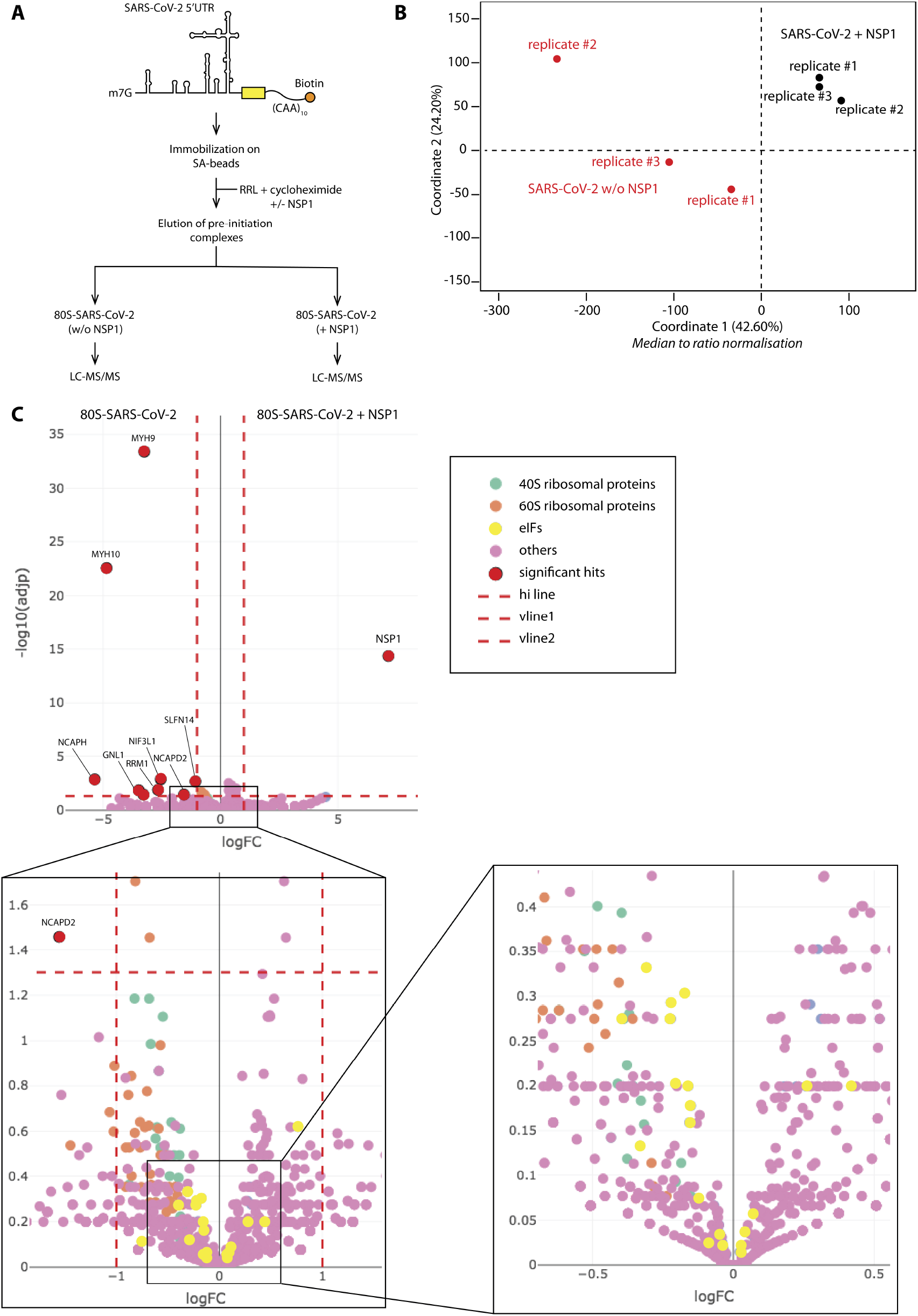
NSP1 remains bound to the pre-initiation complex programmed with SARS-CoV-2 5’UTR. (A) Experimental strategy adapted from (Chicher et al., 2015) to purify translation initiation complexes programmed with SARS-CoV-2 5’UTR in absence or in the presence of recombinant NSP1 (B) Multidimensional scaling plot illustrating global variance and similarities between the SARS-CoV-2 (w/oNSP1) and SARS-CoV-2 (+NSP1) populations detected in the replicates, after a median-to-ratio normalization. (C) Volcano plot showing the proteins co-purified with NSP1 as compared to the control condition performed without NSP1. *Y*- and *X*-axis display adjusted *p*-values and fold changes, respectively. The proteins indicated by a red circle are enriched in either the SARS-CoV-2 (+NSP1) condition (Log2FC>1) or in the SARS-CoV-2 (w/oNSP1) condition (Log2FC<-1). The dashed line indicates the threshold above which proteins are significantly enriched (adjP < 0.05). Green and orange circles label the 40S and 60S ribosomal proteins respectively, yellow circles label the initiation factors, and purple circles correspond to other proteins. The source data are available in Table 2.

## Discussion

Altogether, our data enable us to propose the following model (Figure 6). During the early stages of the SARS-CoV-2 infectious program, ORF1a is translated by canonical cap-dependent translation. The corresponding protein is then processed into NSP proteins. Among these, NSP1 binds to the ribosome with a high affinity. Its C-terminal domain interacts with the mRNA channel entry site and thereby blocks the access to mRNAs. Cellular translation is consequently drastically shut down. However, viral translation escapes this blockage and still goes on. We have confirmed that viral translation is proceeding further in the presence of NSP1 in a so-called viral evasion. We demonstrated here that SL1 is required for this viral evasion. Deletion of SL1 abrogates NSP1 evasion which is in good agreement with a previously published model in which viral translation is also inhibited by NSP1 (Schubert et al., 2020). Indeed, according to the primers described in this publication, the reporter used in the later study did not contain SL1, which in fact confirms our results obtained with Δ40, Δ60, Δ80 and Δ150 (supplemental Figure 1) that showed that viral evasion is abrogated when SL1 is deleted.

**Figure 6:**
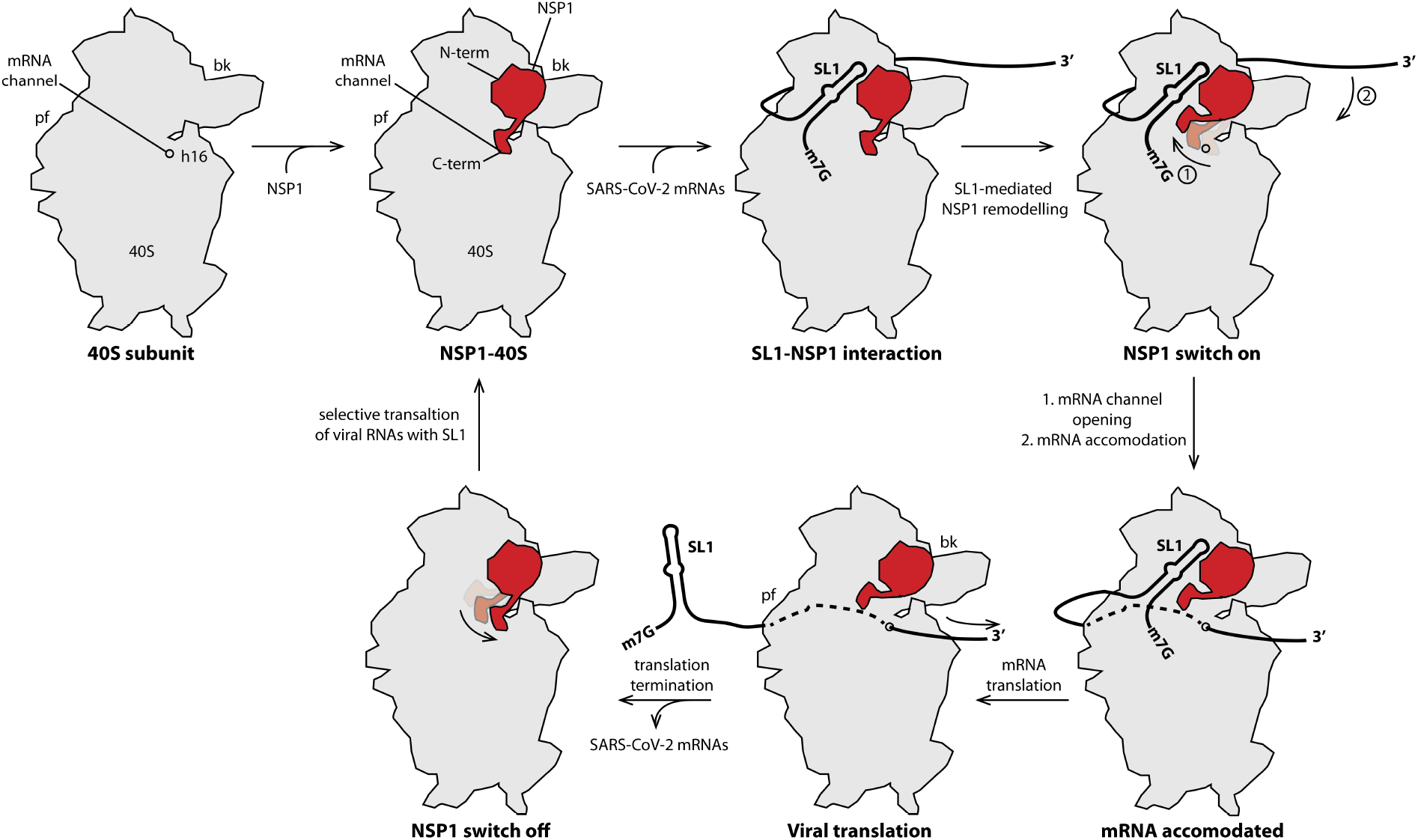
Model for NSP1 acting as a gate-keeper to ensure NSP1 evasion by SARS-CoV-2 5’UTR. In the early phase of infection, NSP1 protein is produced and binds with high affinity to the 40S ribosomal subunit. The cartoons of the 40S subunits are shown from the solvent side. NSP1 binds on the beak of the 40S by interacting with the N-terminal part of NSP1, this interaction places the C-terminal part of NSP1 in the mRNA channel entry site and thereby prevents any mRNA accommodation. The viral mRNA transcripts contain in their 5’UTR SL1 that by interacting with the 40S-NSP1 complex, enables mRNA accommodation and the formation of translation initiation complexes (the mRNA is accommodated in the decoding site on the 40S intersubunit side and is shown with dashed line). This required the removal of the C-terminal domain of NSP1 to open the access to the mRNA channel. Then, translation initiation, elongation and termination can proceed further. After termination, the mRNA is released, NSP1 C-terminal can refold in front of the mRNA channel and prevents any de novo cellular mRNA translation. Only viral mRNA transcripts can access to the mRNA channel, thanks to SL1 that is present in the 5’UTR of genomic and sub-genomic RNAs.

In the model deduced from the available and the present data (Figure 6), the SARS-CoV-2 5’UTR contains a *cis*-acting element, the SL1 hairpin, that induces during translation initiation a structural rearrangement of NSP1, especially of its C-terminal domain. This frees the access to the mRNA channel and allows viral translation to proceed further. This model is in good agreement with two models recently proposed by others (Banerjee et al., 2020; Shi et al., 2020). They found that the N-terminal part of NSP1 is required for evasion from NSP1 inhibition, that the length of the linker between the N- and C-terminal domains is critical and that the 5’UTR is required for viral evasion. In addition, they proposed that NSP1 is removed from the ribosome and retained by an interaction between the N-terminal domain of NSP1 and the 5’UTR (Banerjee et al., 2020; Shi et al., 2020). Although we cannot rule out that this possibility, the fact that we have not been able to detect any RNA binding ability with free NSP1 is a strong argument against this suggestion. Moreover, if NSP1 is released from the ribosome, this will lead to free ribosomes that can start again to translate cellular mRNAs. To ensure efficient shut down of host translation, it is more efficient for the virus to maintain NSP1 on the ribosome. Therefore, we rather suggest that NSP1 stays bound on the ribosome. Within this model, at the end of translation, the C-terminal domain of NSP1 folds back into the mRNA channel and prevents any *de novo* cellular translation. In the late phase of infection, nine sub-genomic RNAs are produced. Interestingly, all these viral transcripts contain the so-called leader body junction that contains SL1, SL2 and SL3 (Kim et al., 2020). The presence of SL1 in all the SARS-CoV-2 transcripts probably ensures efficient NSP1 evasion while still allowing efficient translation. This is especially important for the translation of sub-genomic RNAs that is required in the late phases of the infectious process when the concentration of NSP1 is high and when most if not all the ribosomes are blocked by NSP1. In the model presented in Figure 6, NSP1 acts as a gatekeeper to control selectively the access to the mRNA channel, preventing cellular translation and restricting translation to the sole viral transcripts. The release of the NSP1 gatekeeper is controlled by SL1 from viral transcripts. In conclusion, the sole presence of SL1 all the SARS-CoV-2 transcripts is a pre-requisite to complete the viral infectious program. Therefore, targeting SL1 for therapeutic purposes would be an elegant approach to impair viral translation, in the early phase but also in the late phases of SARS-CoV-2 infection.

Targeting NSP1 for the development of novel therapeutic strategies has been proposed earlier (Huang et al., 2011a; Jauregui et al., 2013; Jimenez-Guardeño et al., 2015; Kamitani et al., 2006, 2009; Lokugamage et al., 2012a; Narayanan et al., 2008; Tanaka et al., 2012; Tohya et al., 2009; Wathelet et al., 2007; Wu et al., 2020; Züst et al., 2007). Another attractive alternative would be to target SL1. Indeed, SL1 being present in all the viral transcripts, drug-design against SL1 would allow to target viral translation in the early phase of infection by impairing genomic RNA translation and in the late phase of infection by blocking translation of sub-genomic RNAs. Altogether, SL1 might be seen as genuine ‘Achille heel’ of SARS-CoV-2.

## Materials and methods

### *In vitro* transcription

The different variants of reporter constructs were transcribed by run-off *in vitro* transcription with T7 RNA polymerase. Uncapped RNAs were separated on denaturing PAGE (4%) and RNA were recovered from the gel slices by electroelution. The resulting pure RNA transcripts were capped at their 5’ end with the ScriptCap m7G Capping System (Epicenter Biotechnologies).

### *In vitro* translation

*In vitro* translation with cell-free translation extracts were performed using self-made rabbit reticulocyte lysates (RRL) as previously described (Martin et al., 2011). Briefly, reactions were incubated at 30 °C for 60 min and included 200 nM of each transcript and 10.8 μCi [^35^S]Met. Aliquots of translation reactions were analyzed for *Renilla* luciferase activity on a luminometer.

### Sucrose gradient analysis

For sucrose-gradient analysis, 5′-^32^P-labeled mRNA were incubated in self-made RRL, in the presence of recombinant NSP1. NSP1 was incubated with RRL 5 min at 30°C prior to addition of radiolabeled mRNAs. The translation initiation complexes were separated on a 10–30% linear sucrose gradient in buffer (25mM Tris–HCl [pH 7.4], 50mM KCl, 5 mM MgCl2, 1 mM DTT). The reactions were loaded on the gradients and spun (23,411xg for 2.5h at 4 °C) in a SW41 rotor. mRNA sedimentation on sucrose gradients was monitored by Cerenkov counting after fractionation.

### Primers used

The primers used in this work are listed in table 1.

**Table 1:** primers used in this study

### Mass spectrometry analysis and data processing

Proteins were digested with sequencing-grade trypsin (Promega) and analyzed by nano LCMS/MS as previously described (Chicher et al., 2015). Digested proteins were then analyzed on a QExactive + mass spectrometer coupled to an EASY-nanoLC-1000 (Thermo Fisher Scientific). MS data were searched against the Rabbit UniProtKB sub-database (release 2020_05, taxon 9986, 43454 sequences) with a decoy strategy. Peptides were identified with Mascot algorithm (version 2.5, Matrix Science) and data were imported into Proline 1.4 software (Bouyssié et al., 2020). Proteins were validated on Mascot pretty rank equal to 1, Mascot score above 25 and 1% FDR on both peptide spectrum matches (PSM score) and protein sets (Protein Set score). The total number of MS/MS fragmentation spectra was used to quantify each protein in each condition performed in three replicates. The statistical analysis based on spectral counts was performed using a homemade R package that calculates fold change and p-values using the quasi-likelihood negative binomial generalized log-linear model implemented in the edgeR package (https://github.com/hzuber67/IPinquiry4). The size factors used to scale samples were calculated according to the DESeq2 normalization method (i.e., median of ratios method). Volcano plots display the adjusted p-values and fold changes in Y and X-axis, respectively, and show the enrichment of proteins in both conditions. P-value were adjusted using Benjamini Hochberg method from stats R package.

### NSP1 overexpression and purification

NSP1 and derivatives (R124A+K125A–Inhibits translation, no mRNA degradation–(Lokugamage et al., 2012a); K164A+H165A—biologically inactive–(Narayanan et al., 2008) were cloned in plasmid pET-His-GST-TEV-LIC-(2GT). This vector overexpresses fusion proteins carrying a 6 His-tag on the N-terminal GST domain followed by TEV protease cleavable site and by the NSP1 native protein or mutants. The fusion proteins were expressed in *E. coli* BL21 Rosetta (DE3) pLysS cells. Cells were grown at 37°C to a cell density of OD_600_ = 0.6. Temperature was decreased to 20°C and cells were induced by addition of 0.1 mM IPTG. Twelve hours after induction, pelleted cells were resuspended in EQ/W buffer (40 mM Na phosphate pH 7.2, 500 mM NaCl, 30 mM imidazole) supplemented with 0.1% Triton X-100, cOmplete^™^ Protease Inhibitor Cocktail (Merck) and incubated on ice for 30 min with 1 mg/mL lysozyme. After lysis by sonification, the cell lysate was centrifuged at 105,000xg for 1h30 and the supernatant was applied to Ni-NTA Superflow resin (QIAGEN) equilibrated in buffer EQ/W. After column washing, NSP1 proteins were eluted from the resin by buffer EQ/W containing 250 mM imidazole. The NSP1 fraction was dialyzed against buffer EQ/W without imidazole overnight. NSP1 fraction was loaded on Glutathione HiCap resin (Qiagen) equilibrated with the dialysis buffer and proteins were eluted by the same buffer supplemented with 50 mM glutathione. The purified 6-His-GST-TEV-NSP1 fusion proteins were subjected to TEV protease cleavage overnight at 4°C (50/1 fusion/TEV molar ratios). NSP1 proteins were separated from the 6 His-GST domain using a last purification step on Ni-NTA resin that retained His-tagged GST and TEV proteins. The pure NSP1 proteins were concentrated, and stored in buffer that contains 50% glycerol at −20°C.

### Electro Mobility Shift Assays (EMSA)

To detect RNA-protein interactions by EMSA, recombinant NSP1 or pure human 40S, 60S and 80S ribosomal fractions were incubated with 50 fmol of 5’ ^32^P-labelled RNA transcripts. Briefly, proteins and RNA were mixed with 20 μg of yeast total tRNA and incubated for 20 min in 10 mM Tris–HCl (pH 7.5), 50 mM KCl, 1 mM DTT, 10% glycerol in 20 μl at 0°C. The RNA-protein complexes were analysed by electrophoresis on native 5% polyacrylamide gels using Tris–50 mM glycine as buffer system and visualized by phosphor imaging.

**Table 2:** Mass spectrometry analysis of proteins in the pre-initiation complexes programmed with SARS-CoV-2 5’UTR in the absence or in the presence of recombinant NSP1.

## Acknowledgments

This work is funded by ‘Agence Nationale pour la Recherche’ (ANR-17-CE12-0025-01, ANR-17-CE11-0024, ANR-20-COVI-0078), by ‘Fondation pour la Recherche Médicale’ (project CoronaIRES), by ‘Fondation Bettencourt Schueller’, by University of Strasbourg and by the ‘Centre National de la Recherche Scientifique’.

